# Integrated Cognitive Assessment: Speed and Accuracy of Visual Processing as a Reliable Proxy to Cognitive Performance

**DOI:** 10.1101/335463

**Authors:** Seyed-Mahdi Khaligh-Razavi, Sina Habibi, Maryam Sadeghi, Haniye Marefat, Mahdiyeh Khanbagi, Seyed Massood Nabavi, Elham Sadeghi, Chris Kalafatis

## Abstract

Various mental disorders are accompanied by some degree of cognitive impairment. Particularly in neurodegenerative disorders, cognitive impairment is the phenotypical hallmark of the disease. Effective, accurate and timely cognitive assessment is key to early diagnosis of this family of mental disorders. Current standard-of-care techniques for cognitive assessment are primarily paper-based, and need to be administered by a healthcare professional; they are additionally language and education-dependent and typically suffer from a learning bias. These tests are thus not ideal for large-scale pro-active cognitive screening and disease progression monitoring. We developed the Integrated Cognitive Assessment (ICA), a 5-minute computerized cognitive assessment tool based on a rapid visual categorization task, in which a series of carefully selected natural images of varied difficulty are presented to participants. Overall 448 participants, across a wide age-range with different levels of education took the ICA test. We compared participants’ ICA test results with a variety of standard pen-and-paper tests that are routinely used to assess cognitive performance. ICA had excellent test-retest reliability, and was significantly correlated with all the reference cognitive tests used here, demonstrating ICA’s ability as one unified test that can assess various cognitive domains.

## 1. Introduction

Brain disorders can cause deficiency in cognitive performance. In particular, in neurodegenerative disorders, cognitive impairment is the phenotypical hallmark of the disease. Neurodegenerative disorders, including Dementia and Alzheimer’s disease, continue to represent a major economic, social and healthcare burden^1^. These diseases remain underdiagnosed or are diagnosed too late ^2^; resulting in less favorable health outcomes. Current routinely used approaches to cognitive assessment, such as the Mini Mental State Examination (MMSE)^3^, Montreal Cognitive Assessment (MoCA)^4^, and Addenbrooke’s Cognitive Examination (ACE)^5^ are primarily paper-based, language and education-dependent and need to be administered by a healthcare professional (e.g. physician). These tests are therefore not ideal tools for wide pro-active screening of cognitive impairment, which can be crucial to earlier diagnosis.

Several studies have emphasized the importance of early diagnosis ^6–8,2,9^ and its role in driving better treatment and improvement of cognition and quality of life ^10^. Therefore, developing new tools for effective, accurate and timely cognitive assessment is key to tackling this family of brain disorders.

Growing attention has been drawn to changes in the visual system in connection with dementia and cognitive impairment ^11–16^. Previous studies have linked visual function abnormalities with Alzheimer’s Disease and other types of cognitive impairment ^17–19^. All parts of the visual system may be affected in Alzheimer’s disease, including the optic nerve, retina, lateral geniculate nucleus (LGN) and the visual cortex ^19^. Therefore, visual dysfunction can predict cognitive deficits in Alzheimer’s Disease ^19,20^.

We therefore developed a rapid visual categorization test that measures subject’s accuracy and response reaction times. These two measures are then summarized to assess participants’ cognitive performance. The proposed integrated cognitive assessment (ICA) test is designed to target cognitive domains and brain areas that are affected in the initial stages of cognitive disorders such as dementia, ideally before the onset of memory symptoms. Thus, as opposed to solely focusing on working memory, the test engages the retina, the visual cortex and the motor cortex, all of them are shown to be affected pre-dementia or in early stages of the disease 2128. The ICA’s focus on speed and accuracy of processing visual information ^29–32^ is in line with latest evidence suggesting that simultaneous object perception deficits are related to reduced visual processing speed in amnestic mild cognitive impairment ^33^. Additionally, the proposed test is self-administered and is intrinsically independent of language and culture, thus making it ideal for large-scale pro-active cognitive screening and cognitive monitoring.

In four different experiments, with a total number of 448 participants, we compared the results of the ICA against a wide range of routinely used standard pen-and-paper cognitive tests. The ICA outcome was significantly correlated with several cognitive domains (e.g. speed of processing, memory, verbal learning, attention, and fluency) as measured by the reference pen-and-paper cognitive tests used here. Overall, we demonstrate that the combination of speed and accuracy of visual processing in a rapid visual categorization task can be used as a reliable measure to assess and monitor individual’s cognitive performance.

## 2. Material and Methods

### 2.1 ICA test description

The ICA test is a rapid visual categorization task with backward masking ^30,34,35^. One hundred natural images (50 animal and 50 non-animal) with various levels of difficulty were presented to the participants. Each image was presented for 100 ms followed by a 20 millisecond inter-stimulus interval (ISI), followed by a dynamic noisy mask (for 250 ms), followed by subject’s categorization into animal vs non-animal (Figure 1). Images were presented at the center of the screen at 7 degree visual angle. For more information about rapid visual categorization tasks refer to Mirzaei et al., (2013).

**Figure 1.**
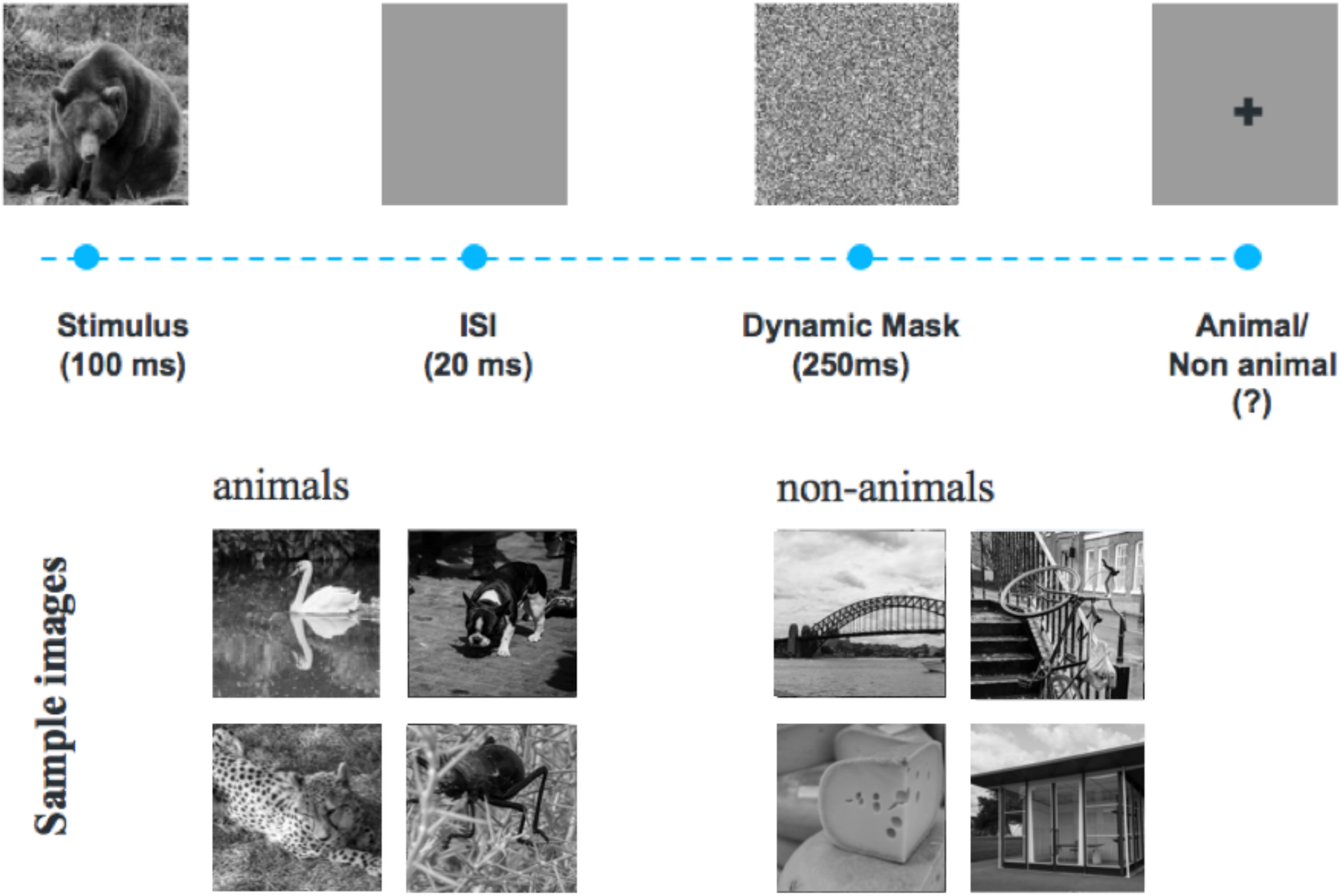
The ICA test pipeline. One hundred natural images (50 animal and 50 non-animal) with various levels of difficulty are presented to the participants. Each image is presented for 100 ms followed by 20 ms inter-stimulus interval (ISI), followed by a dynamic noisy mask (for 250 ms), followed by subject’s categorization into animal vs. non-animal. Few sample images are shown for demonstration purposes.

The ICA test starts with a different set of 10 test images (5 animal, 5 non-animal) to familiarize participants with the task. These images are later removed from further analysis. If participants perform above chance (>50%) on these 10 images, they will continue to the main task. If they perform at chance level (or below), the test instructions will be presented again, and a new set of 10 introductory images will follow. If they perform above chance in this second attempt, they will progress to the main task. If they perform below chance for the second time the test will be aborted.

### 2.2 Scientific rationale behind the ICA test

The ICA test takes advantage of millions of years of human evolution - the human brain’s strong reaction to animal stimuli ^36–39^. Human observers are very good at recognising whether briefly flashed novel images contain the image of an animal, and previous work has shown that the underlying visual processing can be performed quickly ^35,40^. The strongest categorical division represented in the human higher level visual cortex (known as inferior temporal cortex) appears to be that between animates and inanimates. Several studies have shown this in human and non-human primates ^35–37,41,42^. Studies also show that on average it takes about 100ms to 120ms for humans to differentiate animate from inanimate stimuli ^43–45^. Following this rationale, in the ICA test, the images are presented for 100 ms followed by a short inter-stimulus-interval (ISI), then followed by a dynamic mask. Shorter periods of ISI can make the animal detection task more difficult and longer periods reduce the potential use for testing purposes as it may not allow for detecting less severe cognitive impairments. The dynamic mask is used to remove (or at least reduce) the effect of recurrent processes in the brain ^46–50^. This makes the task more challenging by reducing the ongoing recurrent neural activity that could boost subject’s performance. This leaves less room for the resilient brain to compensate for the subtle ongoing neurodegeneration in early stages of the disease.

### 2.3 Participants

As shown in Table 1, we conducted four different experiments; in total, 448 volunteers took part in this study. The study was conducted according to the Declaration of Helsinki and approved by the local ethics committee at Royan Institute. Informed consent was obtained from all participants.

**Table 1.** Summary of all the experiments

| Experiment | Number of Participants | Age mean years, SD [min, max] | Education mean years, SD [min, max] | Gender (#female) | Cognitive Tests | ICA Platform |
| --- | --- | --- | --- | --- | --- | --- |
| 1 | 212 | 74, 10, [46, 98] | 9, 6, [0, 23] | 110 (51%) | MoCA, ICA | Raspberry Pi (RaPi) |
| 2 | 58 | 62, 6, [54, 79] | 14, 5, [3, 24] | 33 (56%) | MoCA, ACE-R, ICA | iPad |
| 3 | 166 | 37, 10, [19, 65] | 14, 3, [1, 20] | 125 (75%) | SDMT, BVMT-R, CVLT-II, ICA | iPad |
| 3' (re-test) | 44 | 38, 12 [18, 64] | 14, 2 [8, 20] | 29 (66%) | ICA, SDMT | iPad |
| 4 | 12 | 29, 3, [20, 36] | 19, 4, [15, 24] | 5 (41%) | ICA | Web |

For each experiment, the table shows number of participants, their demographics (age, education and gender), and the cognitive tests they have taken in each experiment. A total number of 448 volunteers took part in these experiments. Experiments 1 and 2 were to establish ICA correlation with standard-of-care cognitive assessment tools for MCI and dementia screening in older adults (i.e. MoCA and ACE-R). The third experiment covers younger individuals (19 to 65 years-old) who have taken ICA along with three other standard cognitive tests, particularly suitable for this age-range. Experiment 3' is a test-retest: 44 volunteers who participated in the third experiment were called back after five weeks (+/-15 days) to take the ICA and SDMT test for the second time. This was to assess ICA test-retest reliability ( $r=0.96$ , $p<10^{-7}$ ). Experiment 4 is the learning-effect experiment in which 12 young university students took the ICA test every other day for two weeks to see whether ICA is free from learning bias and suitable for micro-monitoring of cognitive performance.

Participants’ inclusion criteria were individuals above age 18, with normal or corrected-to-normal vision, without severe upper limb arthropathy or motor problems that could prevent them from completing the tests independently. For each participant, information about age, education and gender was also collected.

### 2.4 Stimulus set

We used a set of 100 grayscale natural images, half of them contained an animal. The images varied in their level of difficulty. In some images the head or body of the animal is clearly visible to the participants, which makes it easier to detect. In other images the animals are further away or otherwise presented in cluttered environments, making them more difficult to detect. Few sample images are shown in Figure 1. Grayscale images were used to remove the possibility of some typical color blindness affecting participants’ results. Furthermore, color images can facilitate animal detection solely based on color, without fully processing the shape of the stimulus. This could have made the task easier and less suitable for detecting less severe cognitive dysfunctions.

To construct the mask, a white noise image was filtered at four different spatial scales, and the resulting images were thresholded to generate high contrast binary patterns. For each spatial scale, four new images were generated by rotating and mirroring the original image. This leaves us with a pool of 16 images. The noisy mask used in the ICA test was a sequence of 8 images, chosen randomly from the pool, with each of the spatial scales to appear twice in the dynamic mask.

### 2.5 Reference pen-and-paper cognitive tests

#### Montreal Cognitive Assessment (MoCA)

MoCA^4^ is a widely used screening tool for detecting cognitive impairment, typically in older adults. The MoCA test is a one-page 30-point test administered in approximately 10 minutes.

#### Mini-Mental State Examination (MMSE)

The MMSE ^3^ test is a 30-point questionnaire that is used in clinical and research settings to measure cognitive impairment. It is commonly used to screen for dementia in older adults; and takes about 10 minutes to administer.

#### Addenbrooke’s Cognitive Examination-Revised (ACE-R)

The ACE ^51,52^ was originally developed at Cambridge Memory Clinic as an extension to the MMSE. ACE-R is a revised version of ACE that includes MMSE score as one its sub-scores. The ACE-R ^5^ assesses five cognitive domains: attention, memory, verbal fluency, language and visuospatial abilities. On average, the test takes about 20 minutes to administer and score.

#### Symbol Digit Modalities Test (SDMT)

The SDMT is designed to assess speed of information processing, and takes about 5 minutes to administer ^53^. A series of nine symbols are presented at the top of a standard sheet of paper, each paired with a single digit. The rest of the page contains symbols with an empty box next to them, in which participants are asked to write down the digit associated with this symbol as quickly as possible. The outcome score is the number of correct matches over a time span of 90 seconds.

#### California Verbal Learning Test -2^nd^ edition (CVLT-II)

The CVLT-II test ^54,55^ begins with the examiner reading a list of 16 words. Participants listen to the list and then report as many of the items as they can recall. After that, the entire list is read again followed by a second attempt at recall. Altogether, there are five learning trials. The final score, which is out of 80, is the summation of all the correct recalls. As in the brief international cognitive assessment for multiple sclerosis (BICAMS) battery ^56^, we only used the learning trials of the CVLT-II, which takes about 10 minutes to administer.

#### Brief Visual Memory Test–Revised (BVMT-R)

The BVMT-R test assesses visuo-spatial memory ^57,58^. In this test, six abstract shapes are presented to the participant for 10 seconds. The display is removed from view and patients are asked to draw the stimuli via pencil on paper manual responses. There are three learning trials, and the primary outcome measure is the total number of points earned over the three learning trials. The test takes about 5 minutes to administer.

### 2.6 Experiments

We conducted four different experiments, as summarized in Table 1. The first three experiments were designed to measure the ICA correlation with a wide range of routinely used reference cognitive tests. The goal was to investigate whether the speed and accuracy of visual processing in a rapid visual categorization task is correlated with subject’s cognitive performance.

In the first and second experiments, we aimed to test ICA’s ability in assessing cognitive performance in older adults. Therefore, we used MoCA and/or ACE-R as reference cognitive tests, both of which are routinely used to screen for mild cognitive impairment (MCI) and dementia in older adults. In the first experiment, 212 volunteers participated; the ICA test was delivered via a Raspberry Pi (RaPi) platform which is a small single-board computer, attached to a keyboard and a LCD monitor; and MoCA was used as the reference cognitive test. For the second experiment, we had 58 participants; the ICA was delivered via iPad, and both MoCA and ACE-R were used as reference tests in this experiment.

The third experiment had SDMT, BVMT-R and CVLT-II as the reference cognitive tests, measuring speed of information processing, visuo-spatial memory and verbal learning, respectively. These three tests together form the BICAMS battery, which requires about 15 to 20 minutes to administer, and is primarily used to detect cognitive dysfunction in younger adults who may suffer from multiple sclerosis (MS). 166 participants took part in this experiment. Forty-four of them were selected for a re-test as part of a second visit to assess ICA test-retest reliability. The ICA was delivered via an iPad platform.

All the pen-and-paper cognitive tests were administered by a healthcare professional. The administration order for ICA vs. reference cognitive tests was at random.

Finally, experiment 4 was designed to study whether the ICA test had a learning bias if taken multiple times in short intervals. Learning bias is defined as the ability to improve your test score by learning the test simply because of several exposures to the test. 12 young volunteers participated in this study. For convenience, the ICA was delivered remotely via a web platform. Participants took the ICA test every other day for two weeks.

### 2.7 Accuracy, speed, and ICA summary score calculations

#### Preprocessing

We used boxplot to remove outlier reaction times, before computing the ICA score. Boxplot is a non-parametric method for describing groups of numerical data through their quartiles; and allows for detection of outliers in the data. Following the boxplot approach,reaction times greater than q3 + w ^*^ (q3 q1) or less than q1 w * (q3 q1) are considered outliers. q1 is the lower quartile, and q3 is the upper quartile of the reaction times. Where “w” is a ‘whisker’ w = 1.5.

***Accuracy*** is simply defined as the number of correct categorisations divided by the total number of images, multiplied by a 100.

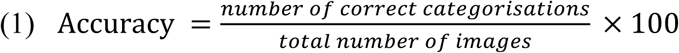

***Speed*** is defined based on participant’s response reaction times in trials they responded correctly.

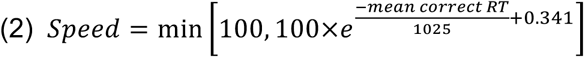

*RT: reaction time*

*e: Euler’s number ∼ 2.7182……*

Speed is inversely related with participants’ reaction times; the higher the speed, the lower the reaction time.

The ***ICA summary score*** is a simple combination of accuracy and speed, defined as follows:

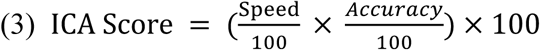

## 3. Results

### 3.1 ICA correlation with the standard-of-care cognitive tests

A key requirement for a clinically useful cognitive assessment test is to establish validity and a correlation with an existing recognized neuropsychological test that is routinely used in clinical practice. Here in three different experiments (see Table 1, experiments 1 to 3), we show that the ICA test is significantly correlated with six standard neuropsychological tests (Figure 2 and Table 2).

**Table 2.** Correlation of the ICA test scores with various domains of cognition.

| Cognitive Test | Correlation with |  |  | Cognitive Domain |
| --- | --- | --- | --- | --- |
|  | ICA | Accuracy | Speed |  |
| **SDMT** | 0.80, $p < 10^{-7}$ | 0.41, $p < 10^{-7}$ | 0.71, $p < 10^{-7}$ | speed of processing |
| **CVLT-II** | 0.66, $p < 10^{-7}$ | 0.40, $p < 10^{-6}$ | 0.56, $p < 10^{-14}$ | verbal learning |
| **BVMT-R** | 0.54, $p < 10^{-6}$ | 0.40, $p < 10^{-8}$ | 0.43, $p < 10^{-7}$ | visual memory |
| **MoCA (1) -RaPi** | 0.46, $p < 10^{-11}$ | 0.48, $p < 10^{-12}$ | 0.19, $p < 0.01$ | General |
| **MoCA (2) -iPad** | 0.55, $p < 10^{-4}$ | 0.56, $p < 10^{-5}$ | 0.19, *ns* | General |
| **ACE-R total score** | 0.60, $p < 10^{-6}$ | 0.52, $p < 10^{-4}$ | 0.28, $p < 0.05$ | total score of 5 domains |
| **ACE \| Attention** | 0.27, $p < 0.05$ | 0.25, *ns* | 0.11, *ns* | attention |
| **ACE \| Memory** | 0.47, $p < 10^{-3}$ | 0.37, $p < 0.01$ | 0.26, $p < 0.05$ | verbal memory |
| **ACE \| Fluency** | 0.45, $p < 10^{-3}$ | 0.25, *ns* | 0.35, $p < 0.01$ | Fluency |
| **ACE \| Language** | 0.53, $p < 10^{-4}$ | 0.62, $p < 10^{-6}$ | 0.12, *ns* | Language |
| **ACE \| Visuospatial** | 0.42, $p < 10^{-3}$ | 0.44, $p < 10^{-3}$ | 0.12, *ns* | visuospatial |
| **MMSE** | 0.33, $p < 0.01$ | 0.36, $p < 0.01$ | 0.08, *ns* | General |

This table shows correlation of the ICA test with a wide variety of standard-of-care tests for cognitive assessment, each measuring various domains of cognition. The two columns for *accuracy* and *speed* indicate the contribution of each of these two components separately toward the ICA score correlation with the reference cognitive tests. The correlations are Pearson correlations, p-values are written in front of each correlation; ns means not-significant ( $p > 0.05$ ).

**Figure 2.**
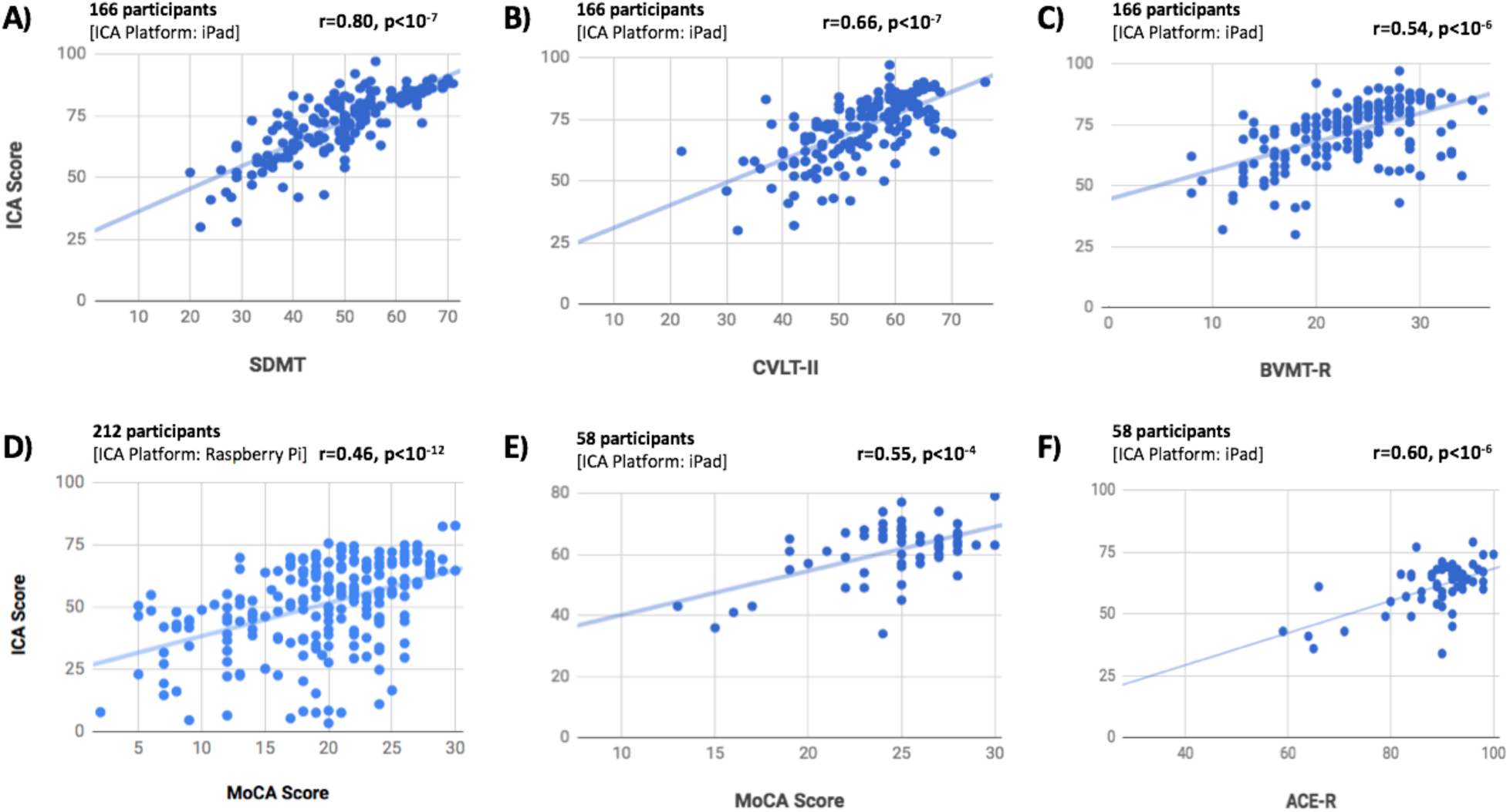
The ICA test score is significantly correlated with a wide range of standard cognitive tests. Participants have taken the ICA test along with one or more standard cognitive tests (see Table 1). Each scatter plot shows the ICA score (y axis) vs. one of the standard cognitive tests (x axis). Each blue dot indicates an individual; the lines are results of linear regression, fitting a linear line to the data in each plot. For each plot, number of participants who have taken the tests and platform on which the ICA is taken are written on top of the scatter plot. ‘r’ and ‘p’ on top-right corner of each plot show the Pearson correlation between the two candidate tests, and the p-value of the correlation, respectively.

**Figure 3.**
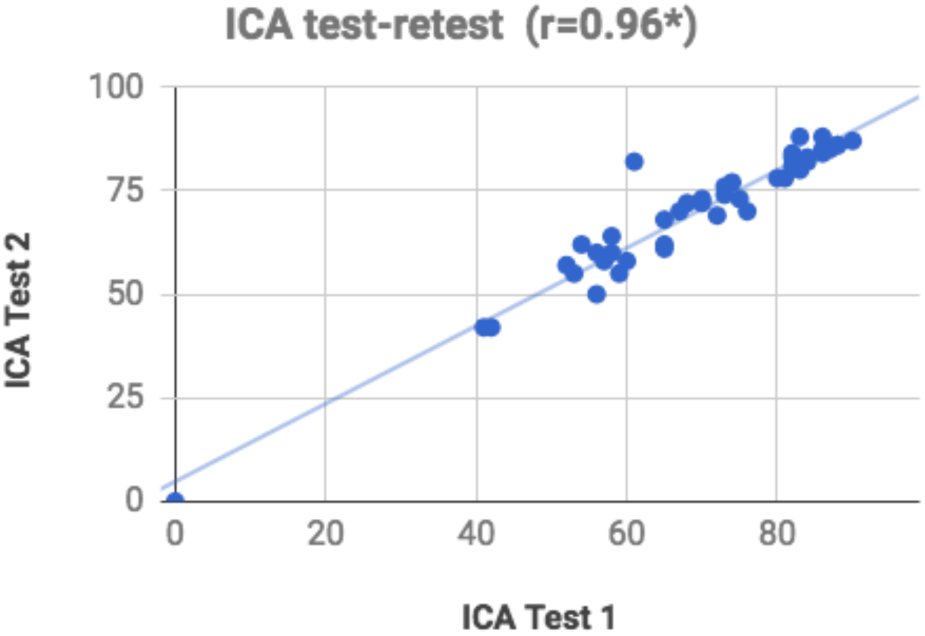
ICA score, test-retest scatter plot. Each blue dot shows the ICA score for an individual taken on two different days. The blue line indicates the linear curve fitted to the test-retest data. [Pearson’s r = 0.96 (*p<10^−7^)]

Given the variability in subject’s demographics, such as age, gender, level of education, etc, a correlation above 0.4 with reference cognitive tests is considered within the acceptable range for construct validity^59^. As an example, cerebral spline fluid (CSF) and blood biomarkers’ correlation with standard cognitive tests, such as MoCA, varies between 0.4 to 0.6 ^60,61^.

We show that the ICA score is significantly correlated with MoCA, tested on two different hardware platforms (RaPi and iPad). ICA correlation with MoCA varies from 0.46 to 0.55 (Figure 2D and 2E) that is within the range for determining construct validity.

The ICA test had a slightly higher correlation with ACE-R (r=0.60, p <10^−6^), compared to MoCA. ACE-R provides a more comprehensive assessment of cognitive abilities and takes a longer time to administer and score (∼ 20 minutes). It is comprised of five subsections, assessing attention, memory, fluency, language, and visuospatial abilities. The ICA correlation with ACE-R (Figure 2F) and its different sub-sections are shown in Table 2. Subject’s MMSE score can also be extracted from the ACE-R test (see Table 2). MoCA and ACE-R are typically used to screen for MCI and dementia in older adults.

In addition, we compared ICA against another set of tests, including SDMT, BVMT-R and CVLT-II (Figure 2A, 2B, 2C, and Table 2) that are more often used in younger individuals to assess cognitive performance. For example, all these three tests are included as part of larger battery of tests that assess cognitive impairment in individuals with MS, such as the ‘minimal assessment of cognitive function in MS’ (MACFIMS) and the ‘brief international cognitive assessment for MS’ (BICAMS).

ICA had the highest correlation with SDMT (r=0.80, p<10^−7^), which is a pen-and-paper test mostly measuring the speed of information processing. CVLT-II, measuring verbal learning, and BVMT-R, measuring visual memory, had correlations of 0.66 and 0.54 with the ICA score, respectively. It is worth noting that a correlation of one is not desirable between the ICA test and any of these cognitive tests, as none of these standard tests are considered the ground truth (or gold standard) in detecting cognitive impairments.

The majority of cognitive tests (Table 2) were more correlated with the accuracy component of the ICA test, except for SDMT and CVLT-II, both of which have got a significantly higher correlation with speed compared to that of accuracy (p<0.001; bootstrap resampling of subjects).

Each reference cognitive test used in this study (shown in Table 2) measured different domains of cognition. The ICA score had significant correlations with all of these tests, suggesting that it can be effectively used as one integrated test to provide insights about different cognitive domains (e.g. speed of processing, memory, verbal learning, attention, and fluency).

### 3.2 ICA shows excellent test-retest reliability

One of the most critical psychometric criteria for the applicability of a test is its reliability. That is the deviation observed when using the same instrument multiple times under similar circumstances.

To assess the reliability of the ICA test, a subgroup of 44 participants from experiment 3 (see Table 1) took the ICA test for the second time after about five weeks (+-15 days). Test-retest reliability was measured by computing the Pearson correlation between the two ICA scores [Pearson’s r = 0.96 (p<10^−7^)]. R values for test-retest correlation are considered adequate if>0.70 and good if >0.80 (Anastasi, 1988).

### 3.3 How much of the ICA score is explained by education?

People with higher levels of education tend to score better in the standard pen-and-paper tests, compared to their age-matched group that fall into the same cognitive category. This makes ‘the level of education’ a confounding factor for cognitive assessment.

We were interested to see how much of the ICA score is explained by education in comparison to other cognitive tests. To this end, we computed the Pearson’s correlation between participants’ test scores and their level of education (in years). Explained variance is defined as the square of this correlation, and indicates how of much of the variance of these test scores can be explained by education (Figure 4).

**Figure 4.**
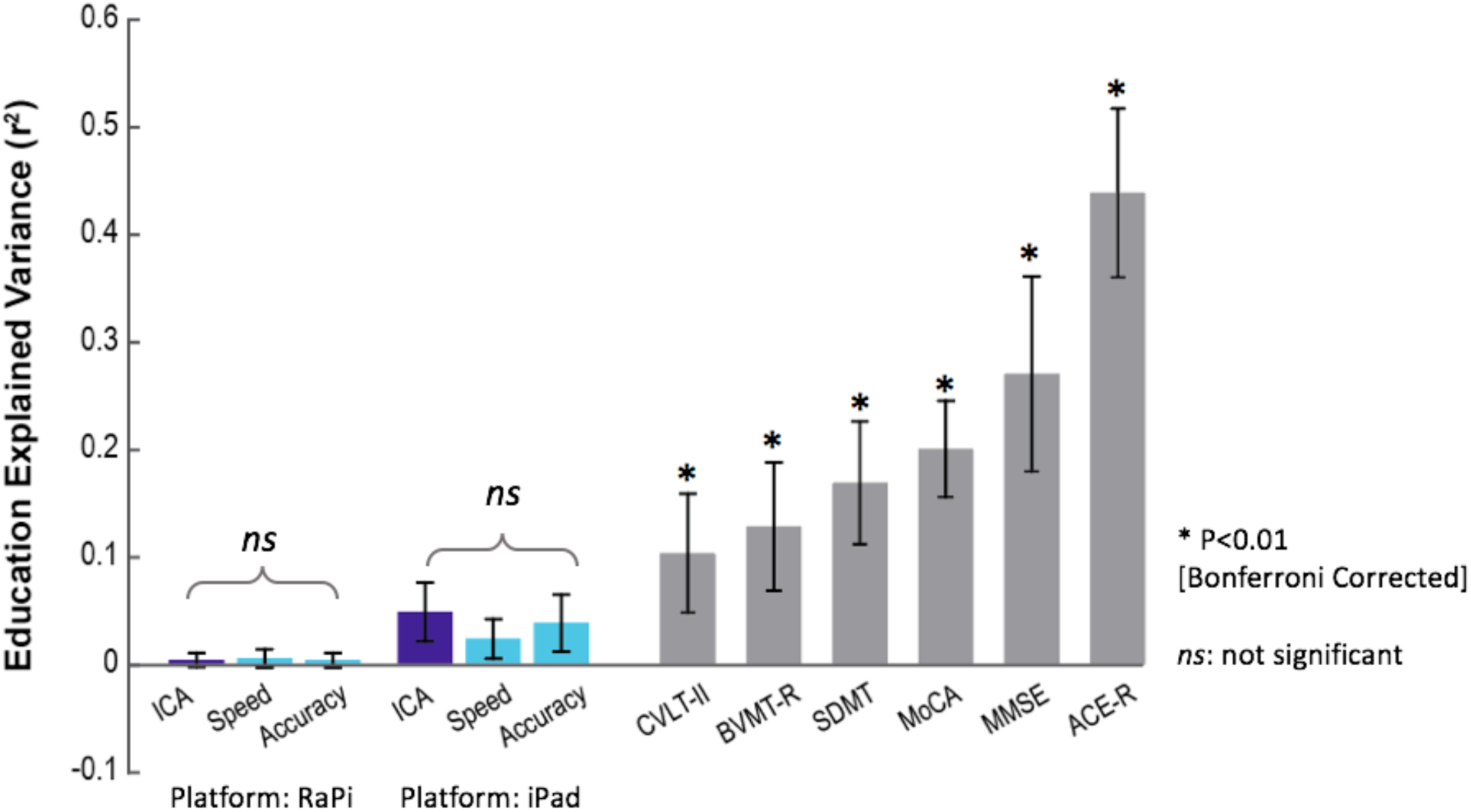
Dependency of standard-of-care cognitive tests on education. Bars indicate how much of the scores reported in each test are explained by education [explained variance = (Pearson’s r)^2^]. Stars show statistical significance, indicating that a significant variance of the test score is explained by education. Statistical significance is obtained by permutation test (10,000 permutations). Error bars are standard errors of the mean (SEM) obtained by 10,000 bootstrap resampling of subjects. P-values smaller than 0.01 (after Bonferroni correction for multiple comparison) are considered significant. ‘ns’ means not significant. The results for ICA (RaPi platform) are based on 212 subjects (Experiment 1 in Table 1); results for ICA (iPad platform) are based on the combined data from Experiments 2 and 3 (224 participants in total) all of whom took the ICA on an iPad. The results for MoCA are based on the combined data from Experiments 1 and 2, in both of which participants took MoCA (270 participants in total). ACE-R and MMSE results are based on the data from Experiment 2. Results for SDMT, BVMT-R and CVLT-II are based on the data from Experiment 3.

We find that a significant variance of all the standard cognitive assessment tests is explained by education, whereas the ICA test does not show a significant relationship with education (Figure 4). In Figure 4, we separately reported the ICA test results for the RaspberryPi platform (Experiment 1: 212 participants) and the iPad platform (experiments 2 and 3: 58 + 166 = 224 participants).

Furthermore, we formally tested whether the ICA score is independent of education using a non-parametric test of independence ^62^. In experiment 1 (i.e. ICA taken on RaPi), the statistical test of independence was positive, showing that ICA score is independent of education (based on 10,000 bootstrap resampling of subjects).

### 3.4 The ICA test has no learning bias

One problem with many existing cognitive tests is that they have a learning bias, meaning that subject’s cognitive performance is improved by repeated exposure to the test as a result of learning the task, without any change in their cognitive ability. A learning bias reduces the reliability of a test if repeatedly used, for example when monitoring performance over time. An ideal test for early diagnosis of cognitive disorders and monitoring cognitive performance would show no ‘learning bias’.

The currently available pen-and-paper tests, such as MoCA, MMSE and Addenbrooke’s Cognitive Examination (ACE), are not appropriate for micro-monitoring of cognitive performance because if identical questions are repeated, healthy participants and those with mild impairment can easily learn the test and improve upon their previous scores as a result of learning rather than any improvement in their cognitive performance.

To investigate whether ICA might be appropriate for such micro-monitoring, we recruited 12 young individuals with high capacity for learning [University students, aged 20 to 36], and asked them to take the test every other day for two weeks (8 days in total). The ICA was delivered remotely via a web platform.

The test data indicate that even in subjects with a high capacity to learn, no learning bias was detected (Figure 5). The ICA score does not increase monotonically, and comparing the mean of the ICA scores across these days, no significant difference was observed (ANOVA, F(7) = 0.62; P-value = 0.73).

**Figure 5.**
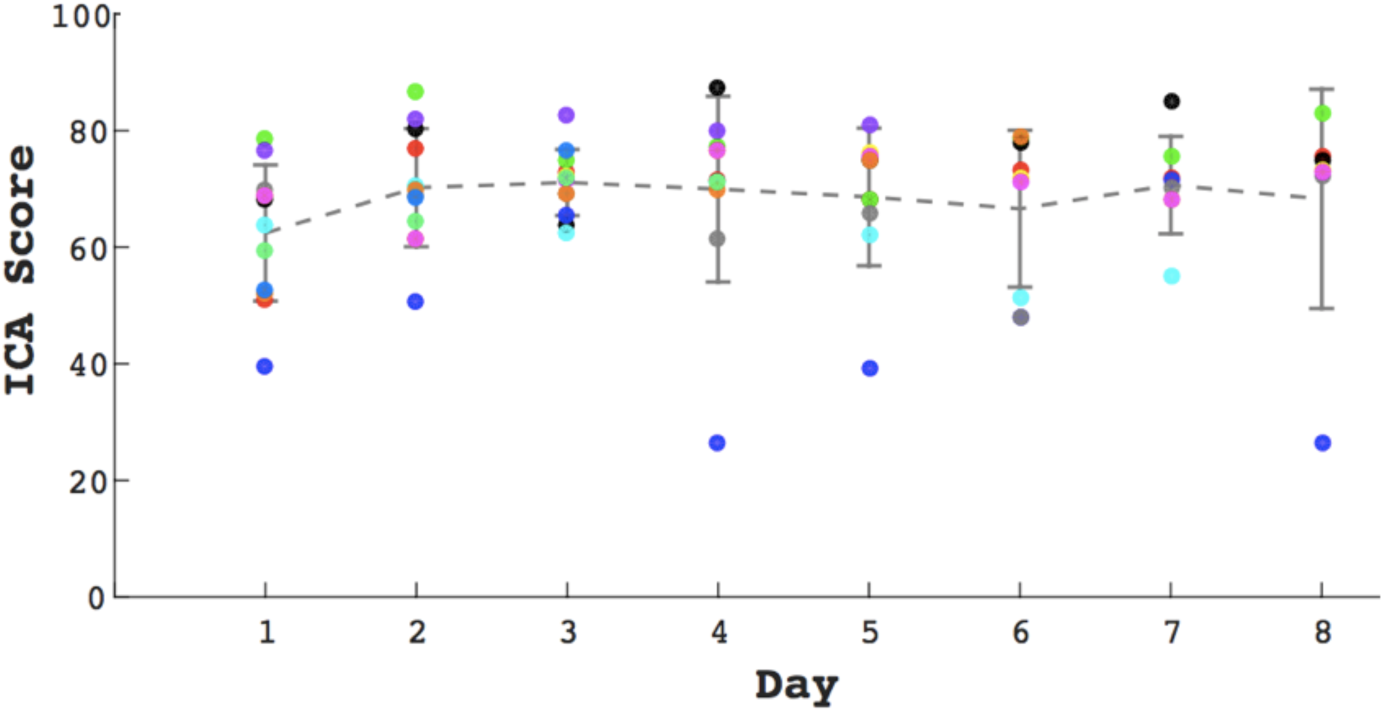
No significant effect of learning in repeated exposure to the ICA test. We find no learning bias when the test is taken multiple times. 12 healthy participants (age range = [20,36]) took the ICA test every other day for over two weeks (ANOVA, F(7) = 0.62; P-value = 0.73). From these 12 participants, 7 of them completed all the sessions (8 days); and the rest did the test for at least the first three days.

## 4. Discussion

Early diagnosis is the mainstay of focus in scientific research ^63,64^. There is currently no available cognitive screening tool that can detect early phenotypical changes prior to the emergence of memory problems and other symptoms of dementia. The vast majority of cognitive tests rely on the patients’ capacity to read and write while more educated individuals can often “second-guess” them. All of these standard tests require a clinician or a health-care professional to administer them, thus adding a considerable cost to the procedure.

We demonstrated that the combination of speed and accuracy of visual processing in a rapid visual categorization task can be used as a reliable measure to assess individual’s cognitive performance. The proposed visual test has significant advantages over the conventional cognitive tests because of its efficient administration, shorter duration, automatic scoring, language and education independency, potential for medical record or research database integration, and the capacity for micro monitoring of cognitive performance given the absence of a “learning bias”. Thus, we suggest ICA as a practical tool for routine screening of cognitive performance.

### 4.1 Potential use of ICA for early detection of dementia

Because of the high compensatory potential of the brain, symptoms of chronic neurodegenerative diseases, such as Alzheimer’s (AD), Parkinson (PD), Huntington (HD) diseases, vascular and frontotemporal (FTD) dementias occur 10–20 years after the beginning of the pathology ^9^. Late stages of these disorders are characterized by massive neuronal death that is irreversible. Therefore, any late therapeutic treatment in the course of the disease will most likely fail to positively affect the disease progression in any meaningful way. This is illustrated by recent failures of anti-AD therapies in late stage clinical trials ^2,65^ Thus further emphasizing the importance of the development of screening tests capable of detecting such diseases in their early asymptomatic stage.

ICA aims at early detection of cognitive dysfunction by targeting brain functionalities that are affected in the initial stages of the neurodegenerative disorders (e.g. dementia), specifically before the onset of memory symptoms. Given the decade-long lag between tissue damage and memory deficits in dementia, the ICA instead examines the visuo-motor pathway. Studies in the past 20 years reveal that all parts of the visual system may be affected in Alzheimer’s Disease, including the optic nerve, retina, lateral geniculate nucleus (LGN) and the visual cortex ^19^. Particularly, in early stages of the disease, brain areas associated with the visuo-motor pathway are affected, beginning with the retina ^22–24,27^, the visual cortex ^21,26,27^ and the motor cortex ^25,28^ so together these represent more effective areas to look for the impact of early stage neurodegeneration as opposed to solely focusing on memory. The ICA focuses on cognitive functions such as speed and accuracy of processing visual information which have been shown to engage a large volume of cortex, while being a predictor of people’s cognitive performance 30–32; thus, monitoring the performance and functionality of these areas altogether can be areliable early indicator of the disease onset.

### 4.2 Suitability for remote and frequent cognitive assessment

Remote monitoring or home-based online assessments is beneficial for patients, clinicians and researchers. Home-based assessment allows for a more comfortable setting for patients with a low stress environment. In addition, researchers and clinicians will have a time-efficient and convenient assessment instrument, which enables a valid and reliable evaluation of individuals’ cognitive performance. Furthermore, online assessment allows the researcher to collect data from a large number of participants in a short time period.

Given that the ICA test is self-administered and that it does not suffer from a learning bias, it can be used remotely and frequently to track changes in individuals’ cognitive performance over time. This makes the test even more useful for early diagnosis, by allowing the test to be used longitudinally, in a design wherein individuals are compared against their own baseline.

## 5. Conclusion and future directions

The ICA is designed to be an extremely easy to use, versatile and practical measurement tool for studies into dementias and other conditions that have an element of cognitive function, as it allows simple, sensitive and repeatable data collection of an overall score of a subject’s cognitive ability. The ICA platform is being further developed to employ artificial intelligence (AI) to improve its predictive power, utilizing patterns of participants’ response reaction times. The AI platform will allow for accurate classification of participants into cognitively healthy or cognitively impaired by comparing their ICA test profile with a large dataset of many individuals with validated clinical status which the AI platform has “learned” from. The AI engine will have the ability to improve its accuracy over time by learning from new data points that are incorporated into its training datasets.

## Acknowledgements

We thank Mark Phillips, and Giulia Paggiola for reviewing the paper prior to submission. We also thank Jonathan El-Sharkawy, Johannes Bausch, and Jean Maillard for help with developing the software needed for data acquisitions. We are also grateful to Mohammad Arbabi for his help with subject recruitment.

## Author contributions

SKR wrote the manuscript. CK gave comments on the manuscript. SKR, SH, MS, HM, MK, SMN, and ES helped with the data acquisitions. SKR, SH, SMN and CK helped with devising the protocol for the experiments. SKR, MS, HM, MK analyzed the data.

## Competing interests

Dr. Khaligh-Razavi, Dr. Habibi, and Dr. Kalafatis serve as directors at Cognetivity ltd. Other authors declared no competing interests.

